# Horizontal transfer of bacteriocin biosynthesis genes requires metabolic adaptation to improve compound production and cellular fitness

**DOI:** 10.1101/2022.07.28.501952

**Authors:** Sophia Krauss, Theresa A. Harbig, Johanna Rapp, Timm Schaefle, Mirita Franz-Wachtel, Leonie Reetz, Ahmed M. A. Elsherbini, Boris Macek, Stephanie Grond, Hannes Link, Kay Nieselt, Bernhard Krismer, Andreas Peschel, Simon Heilbronner

## Abstract

Biosynthetic gene clusters (BGCs) encoding the production of bacteriocins are widespread amongst bacterial isolates and are important genetic determinants of competitive fitness within a given habitat. Staphylococci produce a tremendous diversity of compounds and the corresponding BGCs are frequently associated with mobile genetic elements, suggesting gain and loss of biosynthetic capacity. Pharmaceutical biology has shown that compound production in heterologous hosts is often challenging and many BGC recipients produce initially low compound amounts or show reduced growth rates. To assess whether transfer of BGCs between closely related *S. aureus* strains can be instantly effective or requires elaborate metabolic adaptation, we investigated the intra species transfer of a BGC encoding the ribosomally synthesized and post-translationally modified peptide (RiPP) micrococcin P1 (MP1). We found that acquisition of the BGC by *S. aureus* RN4220 enabled immediate MP1 production but also imposed a metabolic burden, which was relieved after prolonged cultivation by adaptive mutation. We used a multiomics approach to study this phenomenon and found adaptive evolution to select for strains with increased activity of the tricarboxylic acid cycle (TCA), which enhanced metabolic fitness and levels of compound production. Metabolome analysis revealed increases of central metabolites including citrate and α-ketoglutarate in the adapted strain, suggesting metabolic adaptation to overcome the BGC-associated growth defects. Our results indicate that BCG acquisition requires genetic and metabolic predispositions allowing the integration of bacteriocin production into the cellular metabolism. Inappropriate metabolic characteristics of recipients can entail physiological burdens, negatively impacting the competitive fitness of recipients within natural bacterial communities.

**Importance:** Human microbiomes are critically associated with human health and disease. Importantly, pathogenic bacteria can hide in human associated communities and can cause disease when the composition of the community becomes dysbalanced. Bacteriocin producing commensals are able to displace pathogens from microbial communities, suggesting that their targeted introduction in human microbiomes might prevent pathogen colonisation and infection. However, in view of future probiotic approaches, strains are needed that produce high levels of bioactive compounds and retain cellular fitness within mixed bacterial communities. Our work offers insights into the metabolic burdens associated with the production of the bacteriocin micrococcin P1 and highlights evolutionary strategies that increase cellular fitness in the context of production. Most likely metabolic adaptations are broadly relevant for bacteriocin producers and need to be considered for the future development of effective microbiome editing strategies.

## Introduction

It is increasingly recognized that biosynthetic gene clusters (BGCs) allowing the production of antibacterial compounds are omnipresent in bacterial communities (1). These antibacterial compounds are frequently referred to as bacteriocins. They can either be produced ribosomally or by non-ribosomal enzymatic systems and are hugely diverse in terms of molecular size and structure (2). In line with their structural diversity bacteriocins have diverse molecular targets and killing mechanisms and therefore exhibit toxicity towards diverse spectra of bacterial species (2, 3). Bacteriocin-producing bacterial lineages have recently gained increasing attention for their potential to displace pathogens from various human body sites thereby preventing infection (4–6).

Gram-positive staphylococci are one example in this regard. The genus comprises human commensals that are predominantly apathogenic, such as *Staphylococcus epidermidis*, *Staphylococcus capitis*, *Staphylococcus lugdunensis* and *Staphylococcus haemolyticus,* but also the frequently invasive pathogen *Staphylococcus aureus*. Many staphylococcal isolates show inhibitory activity against a diverse range of human nasal commensals and pathogens (7–9). Interestingly, most staphylococcal BGCs appear to be associated with mobile genetic elements such as plasmids, transposons, IS-elements, or chromosomal islands with G+C contents diverging from the genome average (1, 4, 7). This suggests that the BGCs are transferred between strains and lineages and create strain rather than species specific antimicrobial properties (1, 10, 11).

Transfer of antibiotic BGC between strains or species represents a natural system for heterologous expression of antibacterial compounds. Besides providing competitive benefits this might be metabolically challenging for the novel hosts. Indeed, transfer of BGCs between classical antibiotic-producing bacterial species (e.g., streptomycetes) often results in the production of limited amounts of compound in heterologous hosts, and adaptive mutations or changes in nutritional supplies are needed to optimise compound production (12–15). The same might be true in the context of naturally occurring transfer of BGCs between staphylococcal strains and species. Acquisition of bacteriocin BGCs places a burden on the recipient cell. The novel genetic material needs to be propagated and, if functionally expressed, it entails production and secretion of high amounts of toxic secondary metabolites. Precursor molecules need to be channelled from primary metabolism and cellular energy levels might consequently be reduced, entailing metabolic costs for the producer (16). Finally, suboptimal producer-immunity against the compound can further entail physiological burdens (17). Accordingly, it seems plausible that BGC acquisition might represent a mixed blessing for bacterial cells. On the one hand, compound production will provide a competitive advantage when susceptible competitors are present. On the other hand, BGC-associated burdens might reduce fitness of the producer and might require adaptive evolution to optimize compound production and fitness. The physiological costs and the mechanisms of integration of bacteriocin synthesis into primary metabolism in staphylococci remain unclear. However, knowledge about this phenomenon is key to understand why BGCs are largely strain specific and not species-wide conserved traits. Future approaches for the use of bacteriocin-producing bacterial strains to displace pathogens from human microbiomes will crucially depend on the availability of “healthy” strains stably producing high levels of antibiotic molecules.

In this work we used multiomic approaches to study the transfer of the naturally occurring plasmid pD4-19 between closely related *S. aureus* strains. This plasmid encodes the BGC for the biosynthesis of the thiopeptide bacteriocin micrococcin P1 (MP1). Transfer of pD4-19 to *S. aureus* RN4220 allowed immediate production of MP1 but caused growth defects. Genome analysis showed that long-term *in vitro* evolution experiments altered the sequence of the citrate synthase-encoding gene to increase translation of this core metabolic enzyme. Metabolome analysis revealed significant changes in the levels of central metabolic molecules including citrate, α-ketoglutarate and several amino acids. Transcriptome analysis showed that adaptive evolution also increases the expression of ribosomal proteins as well as of enzymes involved in cofactor biosynthesis and protein turnover, all suggesting increased cellular fitness in the context of augmented MP1 production. Phenotypically, the adaptation enhanced compound production and allowed overcoming BGC-associated growth defects. Our data indicate that strain-specific genetic and metabolic predispositions will determine the levels of bacteriocin production and fitness of BGC-recipients. Hence, this will most likely determine the success of a BGC recipient in the context of competitive environments.

## Results

### MP1 BGC as a model system

We sought to investigate the effects of BGC-acquisition on the cellular fitness of the recipient cell. Screening of our extensive collection of nearly 1500 nasal isolates of diverse bacterial species resulted in the identification of two *S. aureus* strains showing intra-species inhibition against the test strain *S. aureus* USA300 LAC. Whole genome sequencing (WGS) and antiSMASH (bacterial version 5.0) analysis revealed the presence of plasmids encoding BGCs for aureocin A70 (18) in strain *S. aureus* P1-22 and for the thiopeptide MP1 (19) in strain *S. aureus* D4-19. The plasmid pD4-19 has a size of 28 391 bp (Fig. 1A) and, besides the 11 kb MP1-encoding BGC (Fig. 1B), encodes a β-lactamase, which we considered useful for *in vitro* plasmid transfer experiments. Therefore, we focused on pD4-19 as a model BGC.

**Figure 1.**
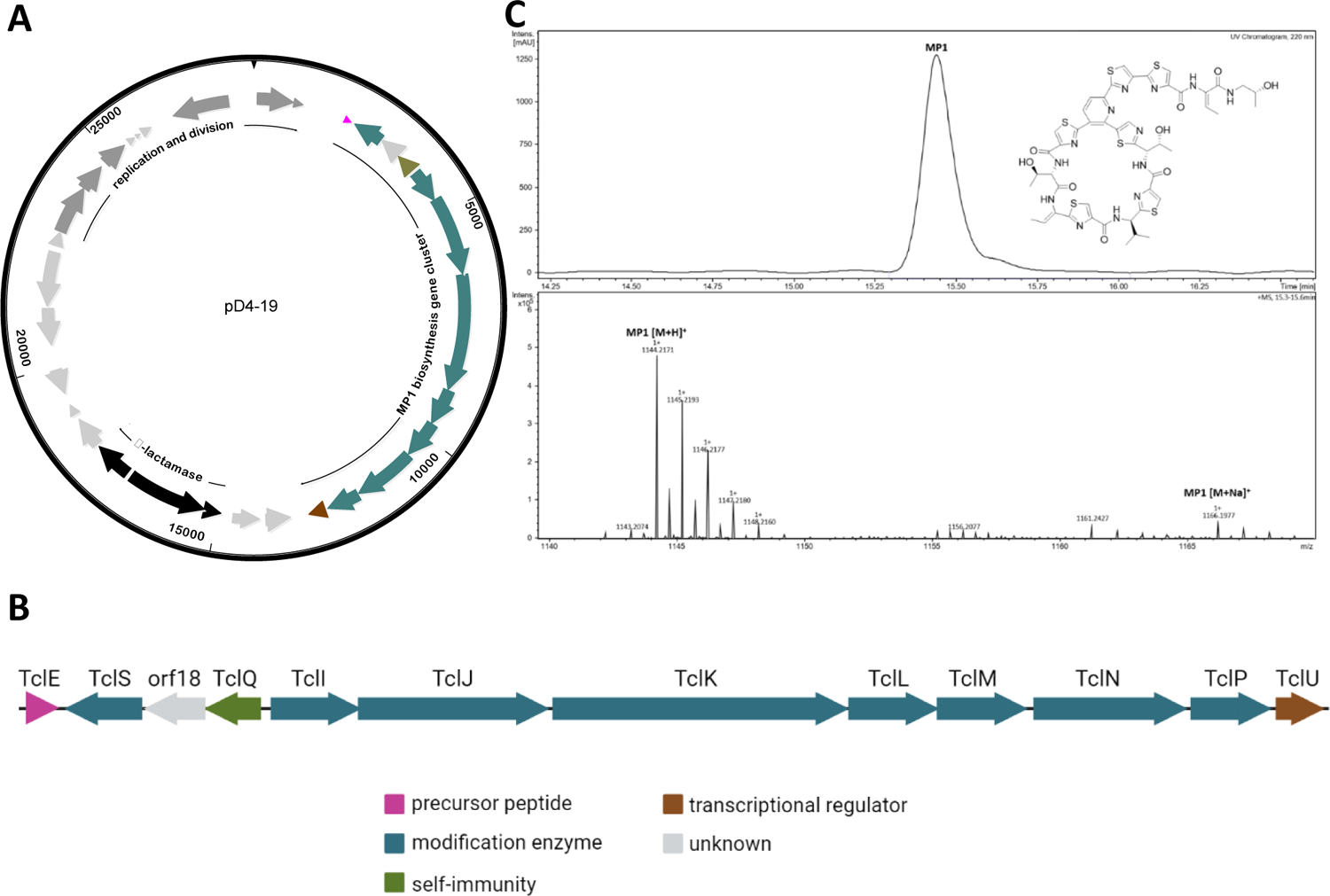
Plasmid pD4-19 encodes the BGC for MP1 production. **A)** Map of plasmid pD4-19 (28 391 bp) that was identified in *S. aureus* D4-19 and that encoded the MP1 BGC (colour), a β-lactamase operon (black), several genes encoding for replication and division factors (dark grey) and hypothetical genes (light grey). **B)** Detailed view of the MP1 BGC. The annotated protein functions are indicated by different colours. Created with BioRender.com. **C)** HPLC-UV-chromatogram (220 nm) of methanol extract from *S. aureus* D4-19 culture pellet. Absorbance was measured in milli absorption units (mAU). The chemical structure of MP1 is depicted. MS spectrum confirming the sum formula of MP1 ([M+H]^+^, m/z = 1144.2171 (found), m/z = 1144.2173 (calculated for C_48_H_50_N_13_O_9_S_6_)).

The thiopeptide MP1 is a RiPP-type (ribosomally synthesized and post-translationally modified peptide) bacteriocin targeting the bacterial ribosome (20), and an alternative ribosomal subunit gene (*tclQ*) provides resistance to the producer (20–22). The MP1 BGC on pD4-19 showed an overall similarity of 99% to the gene cluster identified by Liu et al. in an *S. hominis* isolate (19), suggesting the production of the same compound. To validate this, we performed whole-cell methanol extraction and analysed the extract by high performance liquid chromatography high resolution electrospray (+)-ionisation mass spectrometry (HPLC HR ESI(+)-MS). This analysis revealed the presence of the mass of MP1 (1144.4 Da with the assigned sum formula of MP1: C_48_H_49_N_13_O_9_S_6_) (Fig. 1C), confirming production of MP1 by *S. aureus* D4-19. To prove that the identified BGC is responsible for the observed antimicrobial activity, we performed random transposon mutagenesis. Strains without antibiotic activity were found to carry the transposon within the MP1 BGC (Fig. S1) confirming that MP1 alone was responsible for the antibiotic activity of *S. aureus* D4-19.

### Acquisition of the MP1 BGC enables MP1 production but imposes a metabolic burden

We sought to investigate whether pD4-19 can be transferred to a related *S. aureus* strain and whether this entails significant physiological changes. The plasmid isolated from *S. aureus* D4-19 was used to transform *S. aureus* RN4220 by electroporation. A recovered transformant (RN-T) showed antimicrobial activity against *S. aureus* USA300 LAC suggesting the production of MP1 (Fig. 2A). However, the zone of inhibition was smaller than that of the original producer *S. aureus* D4-19. HPLC-MS and NMR-analysis of cell extracts showed that RN-T produced MP1 as well as a putative derivative with a mass of 1163.2116 Da (Fig. 2B, S2, S3, S4). However, the combined amount of MP1 and the putative derivative was threefold lower than the amount of MP1 produced by *S. aureus* D4-19 (Fig. 2C).

**Figure 2.**
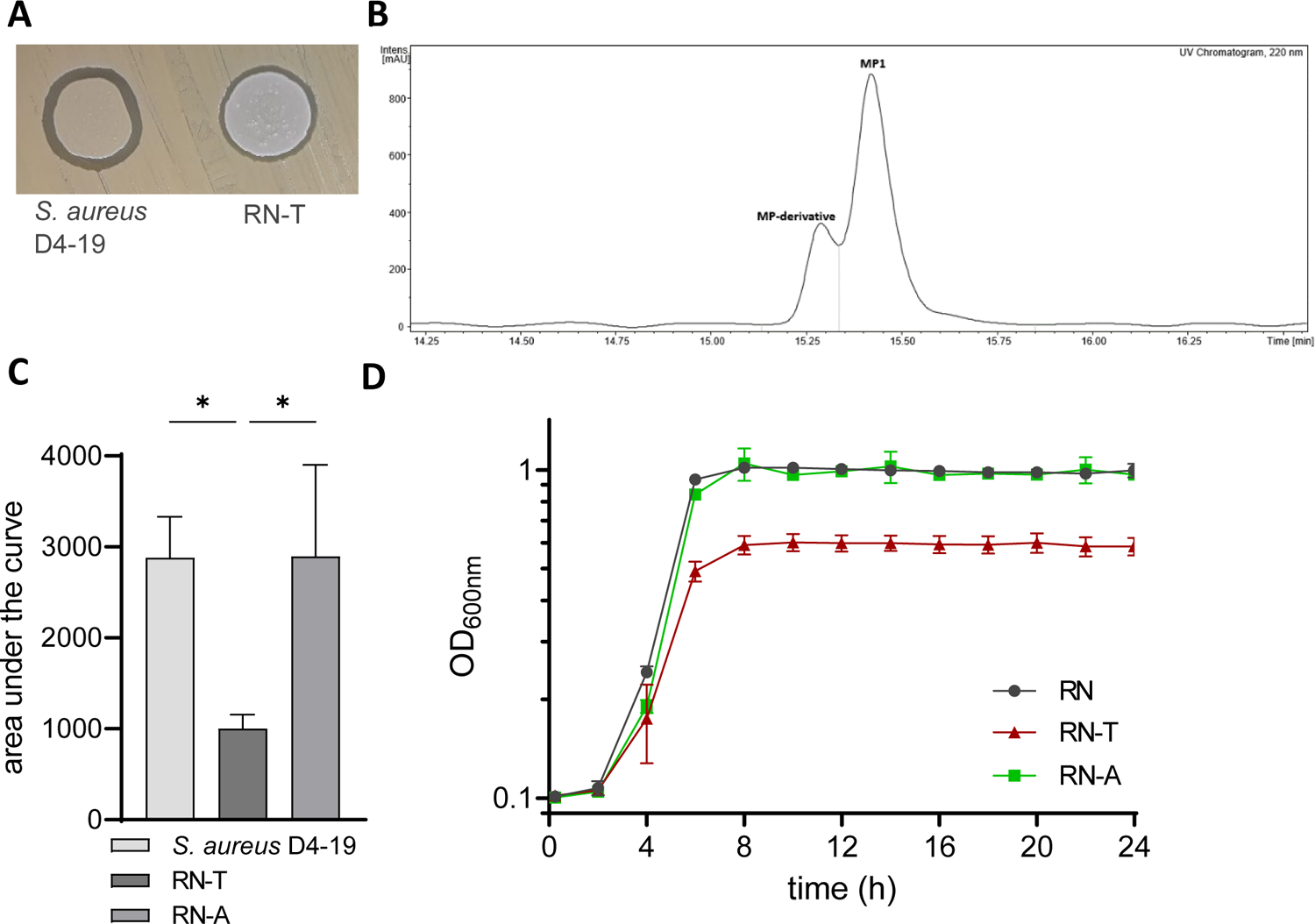
Acquisition of BGC allows production of MP1 but imposes a fitness cost which can be overcome through adaptive evolution. **A)** Spot assay of the nasal isolate *S. aureus* D4-19 and the *S. aureus* recipient strain RN-T on *S. aureus* USA300 LAC, demonstrating antimicrobial activity of both strains against the indicator strain. **B)** HPLC-UV-chromatogram (220 nm) of cell pellet extracts of RN-T, confirming the production of MP1. An MP-derivative (exact mass m/z = 1163.2116), with retention time 15.3 min, is also produced by this strain. **C)** Estimation of the relative quantity of MP1 produced by *S. aureus* strains D4-19, RN-T and RN-A. MP1 amount is estimated as curve integral (n=3). The adapted strain RN-A produces similar amounts as the nasal isolate. MP1 production is significantly reduced in the newly transformed strain RN-T. **D)** Growth curves of *S. aureus* RN, RN-T and RN-A grown in BM over 24 hours in a 24 well plate (n=4). Statistical significance was determined using an ordinary One-way ANOVA (Tukey’s multiple comparisons test) (*p < 0.05).

**Figure 3.**
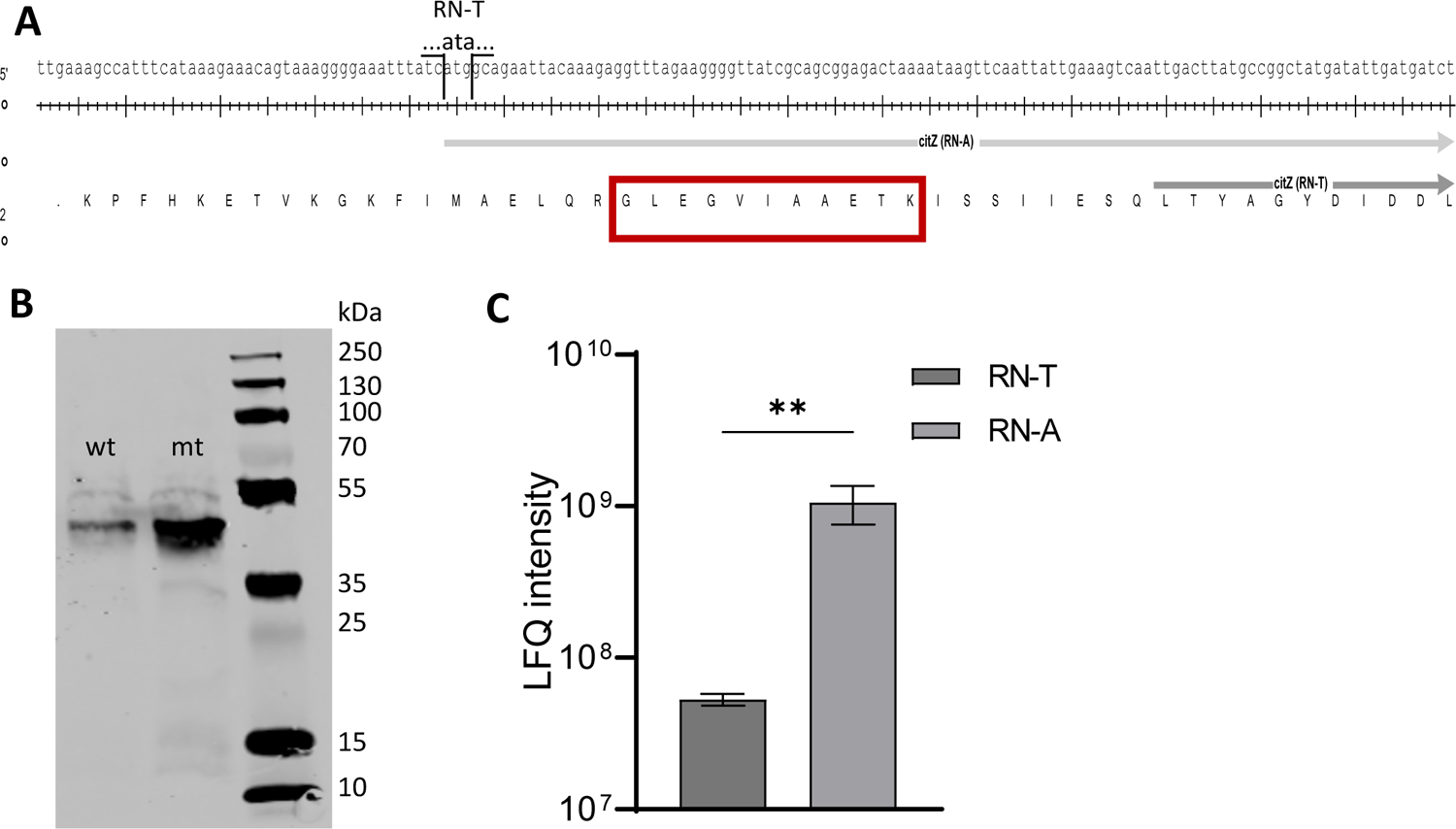
An adaptive mutation in the *citZ* gene increases the levels of the citrate synthase CitZ. **A)** Promoter region and start site of *citZ* (ACCFDFCE_01589) gene from RN-T (wt) (dark grey) and RN-A (mt) (light grey). The wt *citZ* has a delayed start codon compared to the mt allele, where ATA is exchanged to ATG, leading to an earlier transcription start of *citZ* in RN-A. This mutation reverts CitZ to a full-size protein, similar to that of other staphylococci. The red box highlights the part of the amino acid sequence of CitZ that was used for the analysis shown in C). **B)** Western blot of cell extracts from *S. aureus* RN4220 pRB473-XylR-6xHis-*citZ* (wt) and *S. aureus* RN4220 pRB473-XylR-6xHis-*citZ* (mt) expressing the wt CitZ protein and the mt CitZ protein, respectively. CitZ has a size of 42.6 kDa and can be detected in both extracts, but in higher amounts in the mt CitZ expressing strain, indicating leaky expression of wt CitZ. **C)** Proteome data were analysed for the presence of the protein fragment GLEGVIAAETK (depicted in A) (n=3) to confirm leaky expression of wt CitZ with ATA as startcodon. The indicated fragment can be found in both proteomes, but with a 20-fold increase in RN-A. Statistical significance was determined using an unpaired t-test (**p < 0.01).

**Figure 4.**
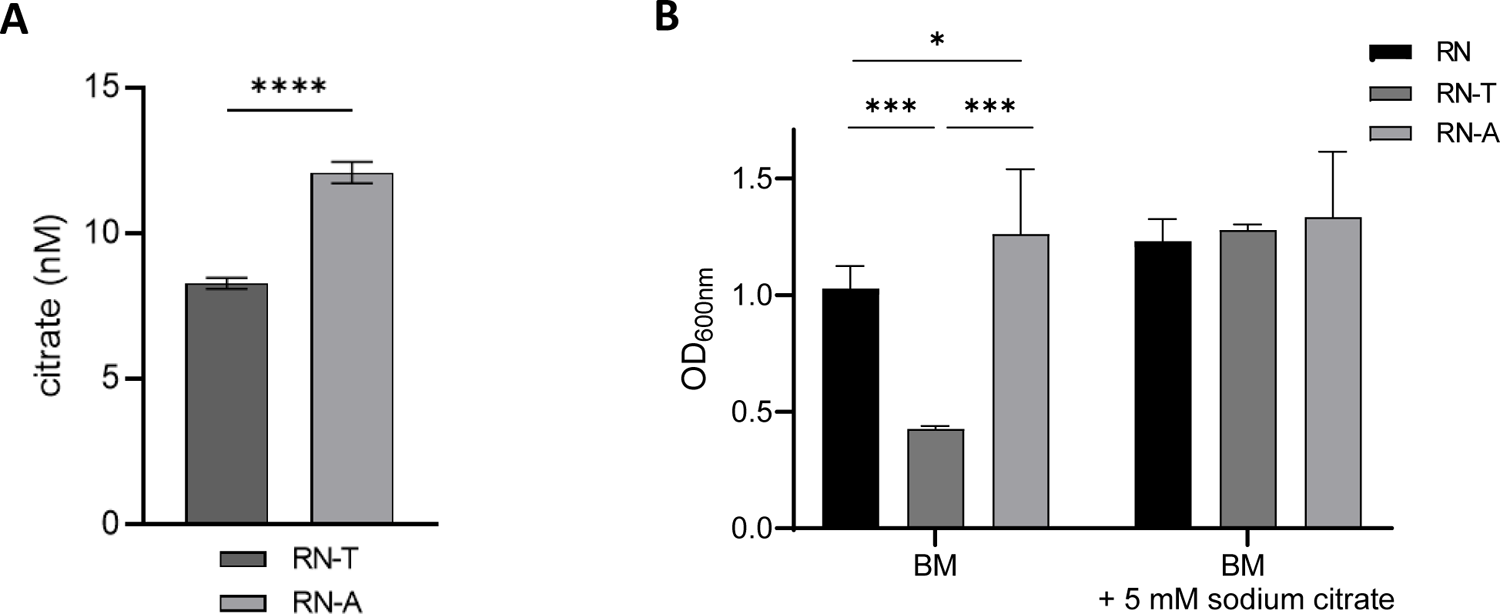
Increased levels of citrate can restore wild type levels of growth in the MP1 BGC recipient strain RN-T. **A)** Significantly reduced citrate levels in RN-T compared to RN-A were determined using the citrate assay kit from Sigma-Aldrich (n=3). Statistical significance was calculated using an unpaired t-test (****p < 0.0001). **B)** Endpoint OD_600nm_ of *S. aureus* RN, RN-T and RN-A in standard growth medium or medium supplemented with 5 mM sodium citrate. Strains were grown in a 24 well plate for 24 hours (n=3). Statistical significance was calculated using an ordinary One-way ANOVA (Tukey’s multiple comparisons test) (*p < 0.05, **p < 0.01, ***p < 0.001, ****p < 0.0001).

Growth curve analysis showed that *S. aureus* RN-T reached lower optical densities than the parental strain *S. aureus* RN4220 (RN) (Fig. 2D) suggesting that MP1 production may limit the proliferation of the strain. To verify that the reduced growth was caused by MP1 production and not by other plasmid-associated effects, a mutant lacking the entire MP1 operon (*tclESQIJKLMNPU* + orf18) was constructed by allelic replacement. The resulting strain *S. aureus* pD4-19 ΔMP1 lacked antimicrobial activity against *S. aureus* USA300 LAC (Fig. S5B) and reached a similar OD_600_ as *S. aureus* RN4220 (Fig. S5A). This supports the idea that acquisition of the MP1 BGC and the associated MP1 production impose a physiological burden on *S. aureus* RN4220. It has been reported that insufficient immunity to an antimicrobial product can limit production of the compound and growth of the producer (17). However, *S. aureus* D4-19 and RN-T mutant strains lacking the MP1 structural gene *tclE* and are therefore deficient in MP1 production were highly resistant to purified MP1 (MIC> 100 µg/ml). This finding indicates that the immunity to MP1 conferred by the alternative ribosomal subunit TclQ is strong enough in both strain backgrounds to allow effective growth even during high-level MP1 production (Fig. S5C).

### Adaptive evolution increases MP1 production and relieves the metabolic burden

We speculated that MP1 biosynthesis might perturb the primary metabolism of *S. aureus* RN4220, thereby limiting bacterial growth and propagation. To analyse whether RN4220 can adapt its metabolism to MP1 production, RN-T was passaged daily in BM medium over 28 consecutive cultures. Colonies arising on solid medium at the end of the experiment were found to be larger than those formed by the original transformant suggesting improved growth. This adapted strain was named RN-A (*S. aureus* RN4220 pD4-19 adapted). Growth curve analysis confirmed that the new strain reached OD_600_ values similar to those of the *S. aureus* RN (Fig. 2D). Additionally, *S. aureus* RN-A produced comparable levels of MP1 as the native MP1 producer *S. aureus* D4-19 and 2.86-fold more than the non-adapted, parental strain RN-T (Fig. 2C).

To identify mutations explaining the phenotypic differences between RN-T and RN-A, the strains were subject to whole genome sequencing and single nucleotide polymorphisms (SNPs) were extracted. We identified a SNP in RN-A, which was located upstream of the gene *citZ* (RN4220 ACCFDFCE_01589) encoding citrate synthase. The mutation created a functional in-frame start codon (ATG instead of ATA) extending the annotated open reading frame of *citZ* by 75 nucleotides (Fig. 3A). An “AG”-rich motif resembling a Shine-Dalgarno sequence is present six base pairs upstream of the new start codon creating a canonical translational start site. In contrast, the annotated shorter allele of *S. aureus* RN4220 lacks an obvious Shine-Dalgarno sequence and relies on a non-canonical “TTG” start codon, which is rarely used in *S. aureus* (23). Accordingly, we speculated that the *citZ* gene of *S. aureus* RN4220 is truncated and potentially non-functional. When the *citZ* allele of *S. aureus* RN4220 was compared with those of other *S. aureus* strains, the shorter *citZ* allele was found in the entire clonal lineage of *S. aureus* RN4220 including the ancestral strains NRS146, NRS133 and VC40 derived from *S. aureus* NCTC8325, which was originally described in 1965 (24). All other 3 834 *S. aureus* genome sequences of the NCBI database, including the original host of the plasmid *S. aureus* D4-19 carry the same full-length *citZ* allele as found in the adapted strain RN-A.

Accordingly, we hypothesised that RN4220 possesses a malfunctioning *citZ* allele whose functionality is restored by the adaptive mutation. To test this, the full-length *citZ* gene of the adapted strain plus its Shine-Dalgarno sequence as well as the corresponding sequence of RN4220 were cloned with a C-terminal His-tag in the *S. aureus* vector pRB473-XylR enabling xylose-inducible expression in RN4220. Upon induction, protein levels were assessed by infra-red Western Blotting. Interestingly, we found that expression of both alleles resulted in the production of apparently full length CitZ proteins (43 kDa) (Fig. 3B). This suggested that the malfunctioning allele is translated using the ATA codon as a non-canonical start codon to produce full length CitZ, a phenomenon that has been described in *E. coli* (25). However, the ATG mutation of the same codon in the adapted allele increased CitZ levels dramatically (Fig. 3B), supporting the idea that the adaptive mutation increases the cellular levels of CitZ by creating an appropriate translational start. This hypothesis was further confirmed by subsequent proteome analysis. An eleven amino acid motif (GLEGVIAAETK) that can only result from the fragmentation of full length CitZ protein was found in both strains. However, this fragment was 20-fold more abundant in RN-A than in RN-T (Fig. 3C). This confirmed that both strains translate the CitZ mRNA using the ATA/ATG codon affected by the point mutation. However, the efficiency of translation is strongly increased by the adaptive mutation entailing increased levels of CitZ.

### Increased citrate levels allow overcoming the MP1-associated burden

To investigate the effects of the mutation on the cellular metabolism, we measured the intracellular concentration of citrate using a colorimetric assay. We found increased levels in RN-A compared to RN-T (Fig. 4A), reflecting the increased amounts of CitZ upon adaptation. Notably, addition of 5 mM sodium citrate to the culture medium allowed the original transformant RN-T to overcome the bacteriocin-associated growth defects (Fig. 4B), indicating that intracellular citrate levels represent the limiting factor for growth of MP1-producing *S. aureus* RN4220.

### The adaptive mutation increases the metabolic activity of the MP1 producer

To gain insights into the overall levels of primary metabolites, we performed Flow Injection Mass Spectrum (FI-MS)-based untargeted metabolomics of cell lysates of *S. aureus* RN, the initial transformant RN-T and the adapted strain RN-A. Hierarchical clustering showed that the metabolic profiles of RN and of RN-A were more similar to each other than to those of RN-T (Fig. 5). This finding indicated that the acquisition of the BGC-encoding plasmid impacts the general metabolism of the recipient, and these effects are largely corrected by the adaptive mutation of *citZ*.

**Figure 5.**
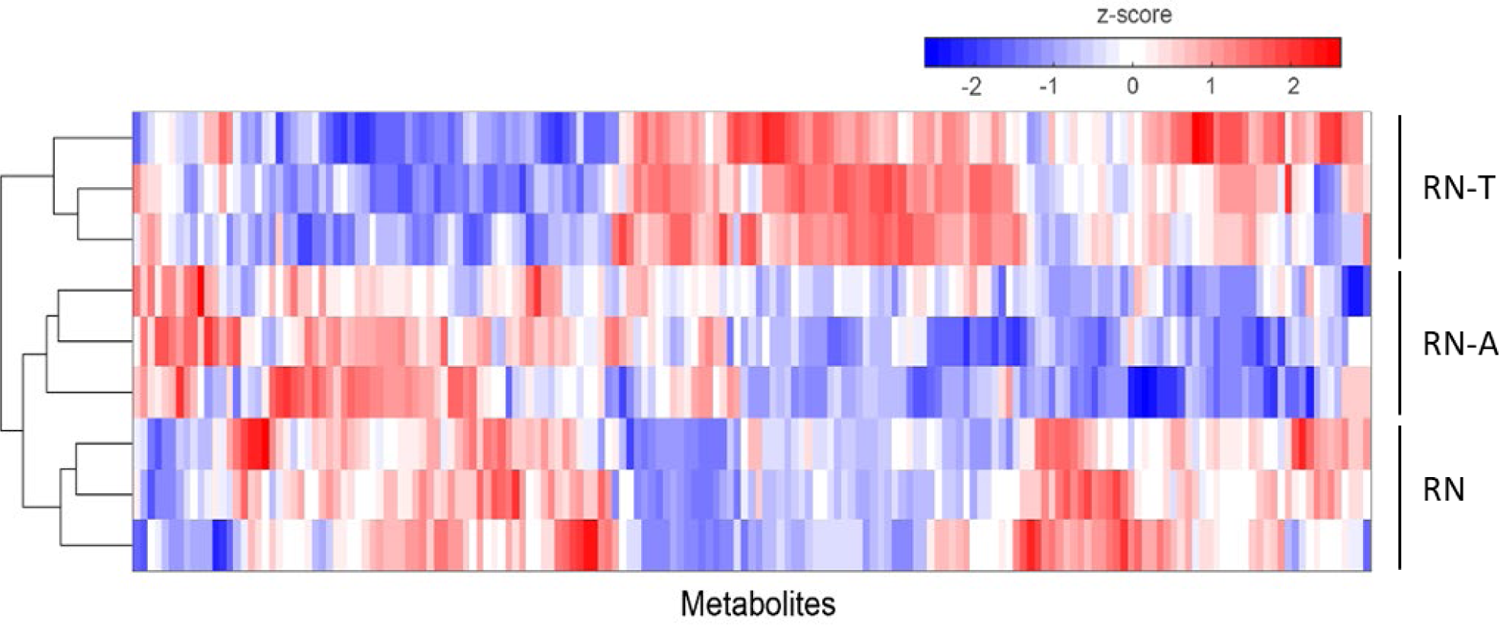
Acquisition of the MP1 BGC alters the metabolite profile of the recipient strain RN-T, an effect that can be reverted by the adaptive mutation. The heatmap shows data from FI-MS analysis for the indicated strains. Depicted are z-score normalized intensities of m/z features in positive and negative ionization mode that were annotated to metabolites (mean of n = 3 replicates). Samples were grouped by hierarchical clustering as indicated by dendrograms.

RiPP-type bacteriocins rely on the availability of appropriate amino acids as well on availability of ATP. Precursors for the biosynthesis of amino acids are derived from intermediates of glycolysis, pentose phosphate pathway and tricarboxylic acid (TCA) cycle. Therefore, we used a combination of LC-MS/MS and FI-MS analysis to investigate the levels of the central metabolites with the highest possible accuracy (Fig. 6, S6, S7, S8, S9). Relative to the WT, the levels of glycolysis intermediates (3-phosphoglycerate, phosphoenolpyruvate, pyruvate, acetyl-CoA) increased upon plasmid acquisition and decreased to or even below RN levels upon adaptive mutation (Fig. 6, S7B). This suggests that the TCA cycle is insufficiently fed upon plasmid acquisition and that the adaptive mutation restores efficient feeding. In line with this, we found increased levels of citrate (citrate and its two isobars isocitrate, 5-dehydro-4-deoxy-D-glucarate cannot be discriminated in this analysis) as well as of α-ketoglutarate upon adaptive mutation (Fig. 6, S7A), supporting the stimulative effect.

**Figure 6.**
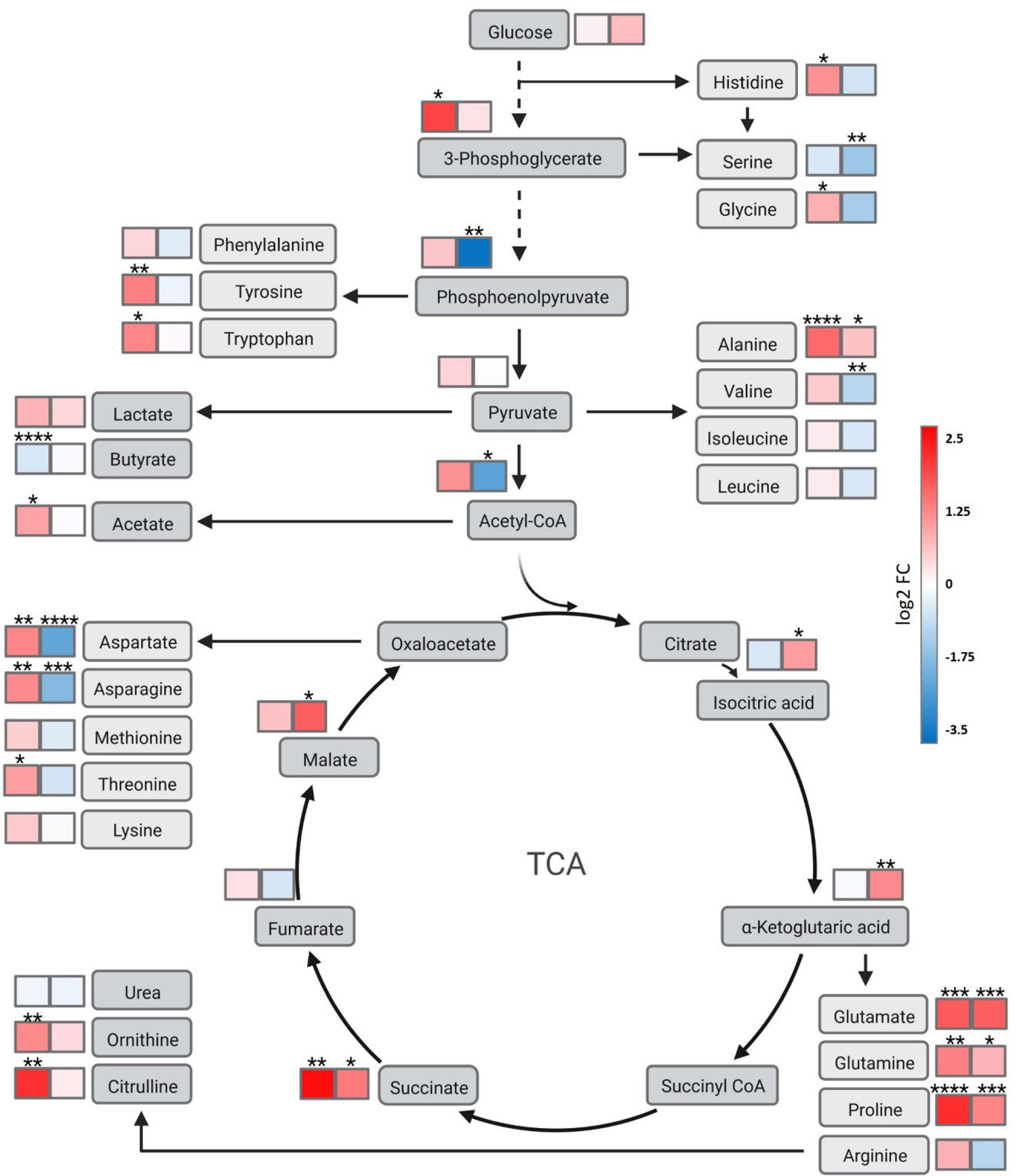
Overview of the metabolic differences between the recipient strain RN-T and the adapted strain. Metabolic differences of RN-T (left box) and RN-A (right box) are shown as log2 fold changes relative to *S. aureus* RN4220. Red: Increase in metabolite levels compared to *S. aureus* RN4220. Blue: Decrease of metabolite levels compared to *S. aureus* RN4220. Metabolic pathways are not depicted in detail. Metabolites were measured via FI-MS and LC-MS/MS. The p-values were calculated using an ordinary One-way ANOVA (Tukey’s multiple comparisons test). Statistical significance is indicated by *p < 0.05, **p < 0.01, ***p < 0.001, ****p < 0.0001. Created with BioRender.com.

Interestingly, we found that pD4-19 acquisition led to the accumulation of various amino acids (aspartate and asparagine, alanine, tyrosine and tryptophan, threonine, glycine, and histidine) pointing to a reduced rate of protein biosynthesis (26). Amino acids did not accumulate in the CitZ adapted strain. In contrast, levels of most amino acids were reduced compared to RN, suggesting their efficient usage in protein biosynthesis (Fig. 6, S6). An exception were glutamate, glutamine and proline, which increased in both RN-T and RN-A relative to RN. Glutamate and glutamine serve as precursors for several amino acids (including proline) and bacteria form glutamate or glutamine by condensation of α-ketoglutaric acid or glutamate with ammonium as a means to acquire environmental nitrogen for anabolic processes. Accordingly, their levels are regarded as an indicator of nitrogen availability (27) and they connect the urea cycle to amino acid biosynthesis. Accumulation of urea cycle intermediates (ornithine, citrulline, argininosuccinate) was only observed in RN-T but not upon adaptive mutation, suggesting that the intermediates are efficiently used to feed the TCA cycle and with it, amino acid biosynthesis in the adapted strain (Fig. 6, S8).

Besides the provision of precursors for metabolic processes, the activity of the TCA cycle is crucial for the aerobic generation of ATP. In the presence of glucose, *S. aureus* produces ATP by substrate-level phosphorylation even in the presence of oxygen and produces predominantly acetate, which is secreted. Only after glucose is depleted, acetate is consumed and decarboxylated using the TCA cycle and the respiratory chain (28, 29). This metabolic switch has been reported to occur after approximately five hours of growth in glucose-containing complex media (29), which matches the observed starting point of growth deficiency in our experiments. We noted that the intracellular levels of acetate were increased in RN-T compared to RN (Fig. 6, S9). We therefore speculated that CitZ and TCA cycle activity might not allow sufficient ATP generation when acetate needs to be catabolized. Interestingly, we detected accumulation of acetate in the culture supernatants of *S. aureus* RN4220 and of RN-T, while accumulation was strongly reduced in RN-A and dropped quickly after six hours of growth (Fig. 7A). This suggests an increased rate of acetate consumption and oxidative decarboxylation upon restoration of the *citZ* allele. However, ATP levels measured by metabolome analysis did not support energy depletion as the reason of the growth defect of RN-T (Fig. 7B). Compared to the WT strain, ATP-levels were only moderately decreased in RN-T and the decrease intensified upon adaptive mutation. As RN-A does not show abnormal growth, these data suggest that even the lowest ATP levels observed in RN-A are sufficient to support maximal growth.

**Figure 7.**
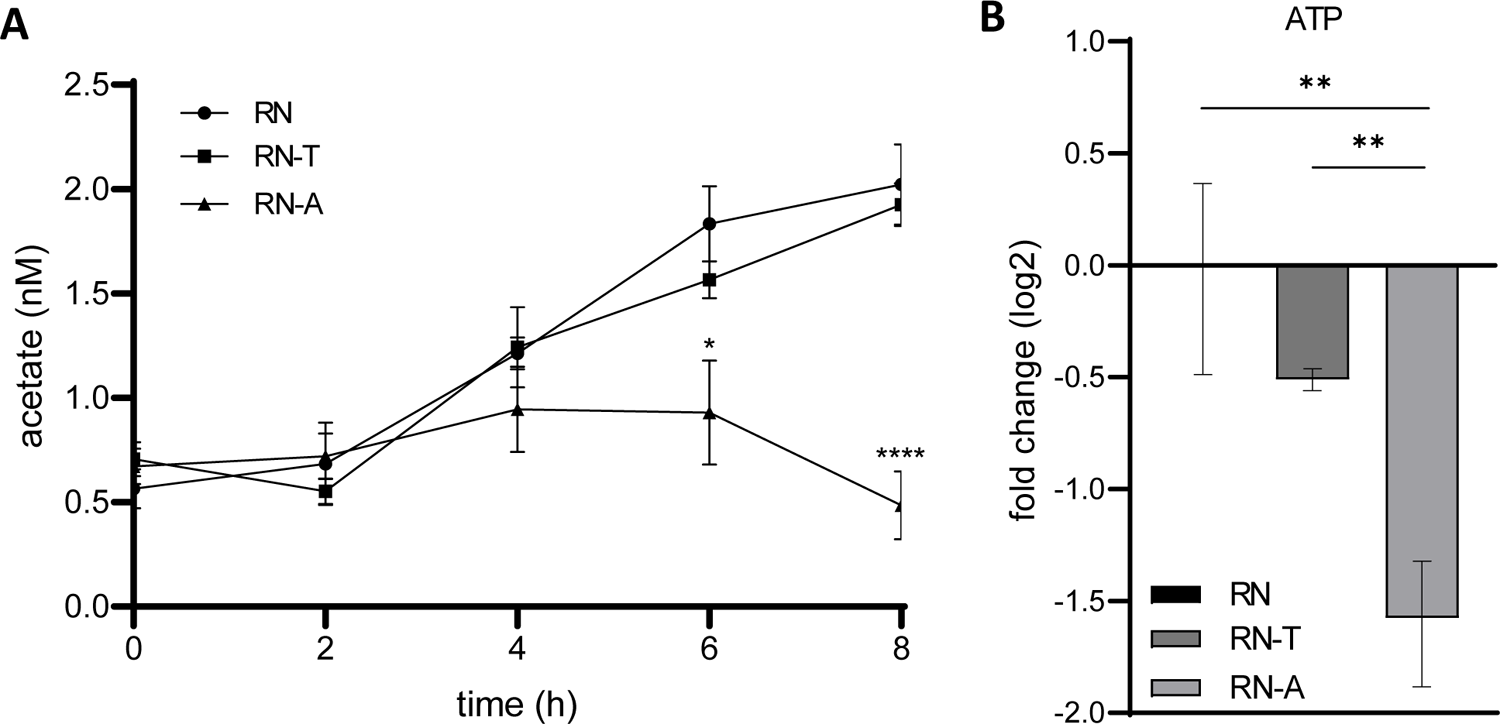
The increased intracellular levels of acetate in RN-T cannot be ascribed to reduced levels of ATP. **A)** Extracellular acetate levels were measured in *S. aureus* RN, RN-T and RN-A after 0, 2, 4, 6 and 6 hours of growth in BM with the acetate assay kit from Sigma-Aldrich (n=3). **B)** Changes (log2-fold) of cytoplasmic ATP levels in *S. aureus* RN-T (dark grey) and RN-A (light grey) compared to RN (black) (n=3). ATP was measured via LC-MS/MS. The p-values were calculated using an ordinary One-way ANOVA (Tukey’s multiple comparisons test). Statistical significance is indicated by *p < 0.05, **p < 0.01, ***p < 0.001, ****p < 0.0001.

### Transcriptomic signatures support increased overall metabolic activity upon adaptive mutation

We performed RNA-seq analysis of RN-T and RN-A to investigate the effects of the adaptive mutation on the transcriptome. We extracted differentially regulated genes and assigned them to Gene Ontology (GO) groups to identify the general cellular functions altered in response to the adaptive mutation in *citZ*. The adaptive mutation entailed increased transcription of the translational machinery (30S and 50S ribosomal subunits), pointing to generally enhanced protein biosynthesis as a consequence of overall metabolic alterations (Fig. 8). This change was accompanied by upregulation of pathways for the biosynthesis of the cofactors folate (*folPBK*), thiamine (*thiEM*), and riboflavin (*ribAB*). Increased amino acid turnover was reflected by upregulation of the L-tryptophan biosynthesis pathway (*trpCDEFG*) as well as of catabolic pathways for threonine (*ilvA*) and alanine (*ald1*). Additionally, upregulation of the genes encoding the urea transporter Utp and of each gene of the urease operon (*ureABCEFGD*), catalysing the hydrolysis of urea into carbon dioxide and ammonia, was observed (Fig. 8, S10A). Furthermore, upon plasmid acquisition, several genes associated with iron homeostasis as well as the genes encoding the twin-arginine translocation (Tat) pathway were differently expressed (Fig. 8). The Tat system is responsible for the translocation of the iron-dependent peroxidase (FepB), which is also involved in iron uptake (30). Accordingly, one can speculate that the increased metabolism upon adaptive mutation does also influence iron homeostasis.

**Figure 8.**
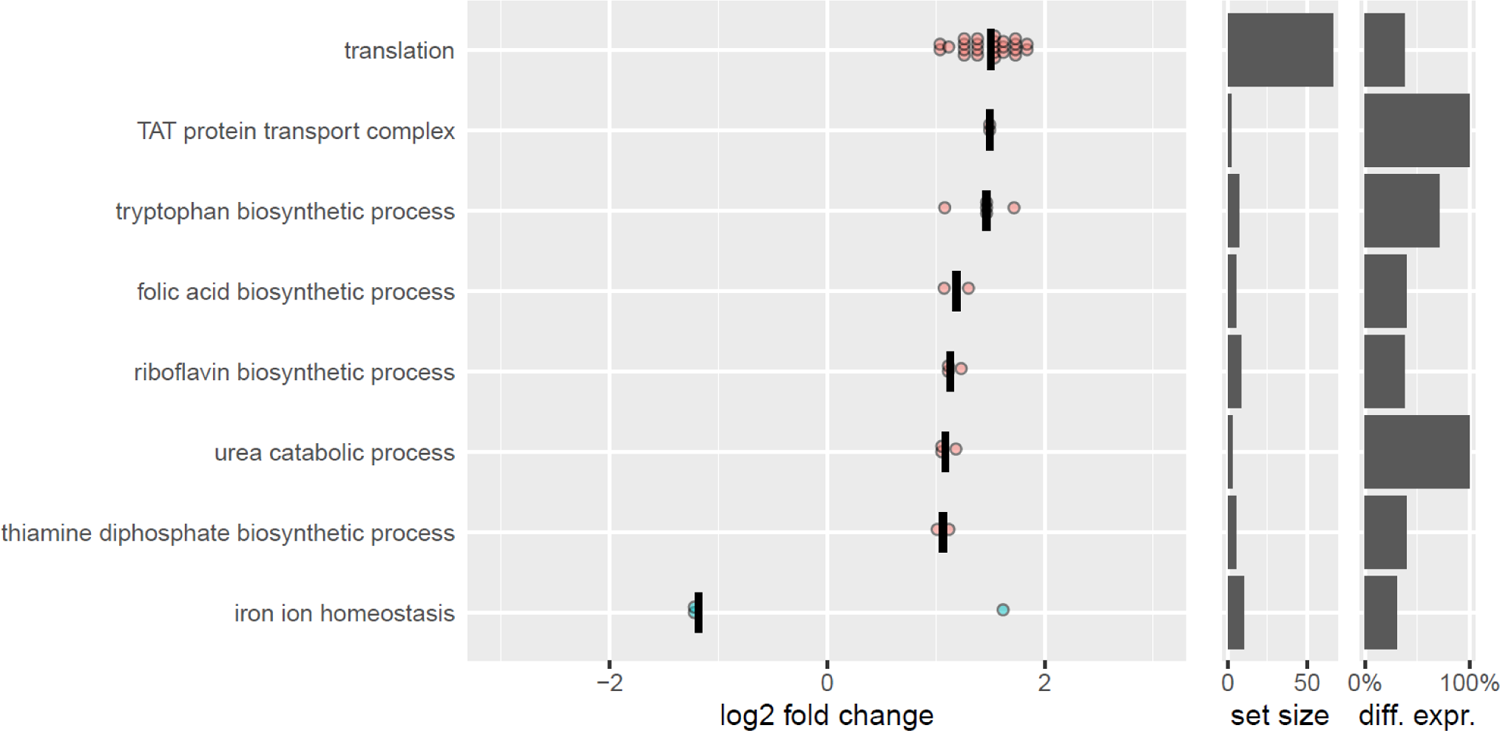
The adaptive mutation increases the overall metabolic activity of strain RN-A as confirmed by transcriptomic analysis. Selection of significantly enriched GO terms and the corresponding differentially expressed genes obtained via RNAseq. Depicted is the log2-fold change in expression levels of *S. aureus* RN-A compared to RN-T. Increased median expression is indicated in red, decreased median expression in blue. The set size indicates the number of genes covered by the GO term. The last column indicates how many genes of the set are differentially expressed (in %).

Of note, expression of plasmid associated genes did not differ significantly between RN-T and RN-A (Fig. S10B).

Inclusion of the transcriptomic profile of *S. aureus* RN into this analysis allowed us to extract several additional effects of plasmid acquisition on the transcriptome. We found transcription of the carnitine transporter *opuC* (*opuCA/CB/CC/CD*) to be downregulated in *S. aureus* RN-T compared to RN. This is in line with reduced carnitine levels detected via metabolome analysis (Fig. S9) (31). Our transcriptome analysis identified several genes that were differentially expressed upon pD4-19 acquisition and whose expression remained altered upon adaptive evolution, arguing for intrinsic effects of the plasmid. Among those genes several were associated with virulence and immune interference including hemolysins (*hlgB/C*, *hlY*), the capsule biosynthesis operon *cap,* the serine protease locus *spl*, and the type seven secretion system *ess* (Fig. S10A). Also transcription of the *icaADBC* operon was increased upon plasmid acquisition and all genes except for *icaD* remained highly expressed upon adaptive mutation. The *ica* operon allows the synthesis of the polysaccharide intercellular adhesin (PIA) and is thereby responsible for biofilm formation (Fig. S10A). In line with this, RN-T showed a significant increase in biofilm formation (Fig. 9). Interestingly, this phenotype was reverted upon adaptive evolution.

**Figure 9.**
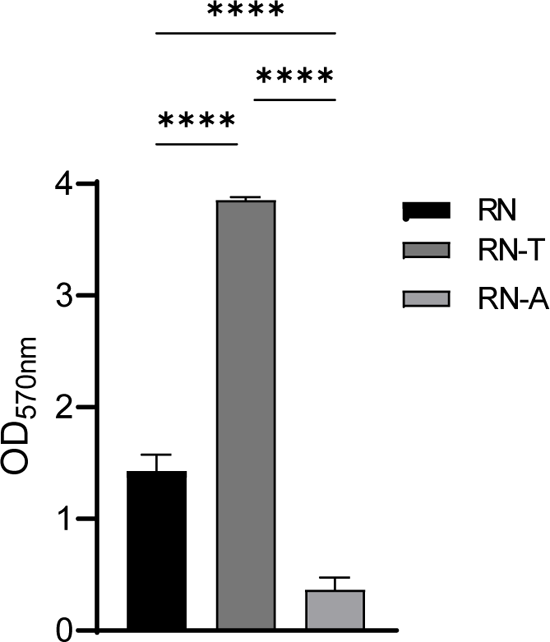
Acquisition of the MP1 BGC is associated with increased levels of biofilm formation. Biofilm formation of *S. aureus* RN, RN-T and RN-A grown in BM (n=3). Statistical significance was calculated using an ordinary One-way ANOVA (Tukey’s multiple comparisons test) (****p < 0.0001).

## Discussion

Using our collection of nasal staphylococcal isolates, we identified the *S. aureus* strain D4-19 carrying a plasmid encoding the MP1 BGC. A number of studies have identified MP1-BGCs in a variety of staphylococcal species, including *S. aureus* (32), *S. epidermidis* (33), *Staphylococcus equorum* (34), *Staphylococcus pseudintermediu*s (35), *Staphylococcus agnetis*, and *S. hominis* (19). Also, an isolate from *Mammaliicoccus sciuri* (former *Staphylococcus sciuri*) (36) has been shown to produce MP1. Moreover, the NCBI database includes various homologous sequences in *S. aureus* genomes as well as many coagulase-negative staphylococci. The abundance of this BGC in different strain backgrounds suggests intra- and inter-species horizontal transfer, which is supported by the fact that the so far identified MP1 gene clusters in staphylococci are all located on plasmids. The cellular consequences of BGC acquisition in staphylococci are profound and we studied the reasons for metabolic constraints along with mechanisms overcoming them using evolutionary adaptation and multiomics approaches.

It is generally recognised that bacteriocin production is associated with metabolic costs for the producing cell and that expression is therefore frequently regulated, often in complex ways for instance depending on producer cell densities or on the detection of bacterial competitors (37, 38). Compound production generates metabolic burdens as the responsible gene cluster needs to be propagated, precursors need to be channelled from the primary metabolism and sufficient self-immunity needs to be established. This concept might be of special relevance for RiPPs such as MP1 as their production relies on cellular tRNA pools and sudden expression upon acquisition of the BGC might therefore disturb central metabolic fluxes. It seems therefore likely that cellular fitness in the context of bacteriocin production requires adjustments of metabolic pathways. Our experiments showed that acquisition of the plasmid pD4-19 enabled production of bioactive MP1. However, the BGC-recipient produced significantly less compound than the original plasmid host (*S. aureus* D4-19) while simultaneously displaying a growth defect compared to the plasmid free parental strain RN4220. Increased availability of citrate, of exogenous or endogenous origin, improved compound production and abrogated any growth defects. Interestingly, we found the recipient *S. aureus* RN4220 to carry a malfunctional *citZ* allele, limiting the levels of citrate synthase activity, of intracellular citrate and most likely, of the entire TCA cycle. Importantly, the entire lineage of *S. aureus* NCTC8325 (parental lineage of *S. aureus* RN4220) developed and maintained the malfunctioning *citZ* allele. Although *S. aureus* RN4220 has been used for decades to study *S. aureus* physiology and virulence, obvious fitness defects compared to environmental isolates have never been reported, suggesting that low-level production of CitZ is in general sufficient for the lineage to thrive efficiently, at least under laboratory conditions. However, the acquisition of pD4-19 made the metabolic shortcomings of the lineage apparent. The most prominent phenotype of TCA cycle-deficient mutants is a growth defect that manifests after approximately five hours of growth in glucose-containing medium (29). This is due to the fact that *S. aureus* preferentially degrades glucose via glycolysis or the pentose phosphate pathway and fermentation to produce acetate (28, 29). Concurrently, catabolite repression of glucose inhibits the TCA cycle (39). Only after glucose is consumed, acetate catabolism demands TCA cycle activity to create ATP and to sustain growth (29). *S. aureus* RN4220 does not show signs of TCA cycle deficiency. However, we observed that acquisition of pD4-19 and the associated MP1 production entailed premature growth arrest as well as increased levels of intracellular pyruvate and acetyl-CoA as well as extracellular accumulation of acetate, all of which are hallmarks of TCA cycle-deficient strains (29). Adaptive mutation reverted these signs of TCA deficiency. Acetyl-CoA levels dropped, and the levels of citrate and α-ketoglutarate increased suggesting efficient feeding of the TCA cycle. Simultaneously, accumulation of extracellular acetate was limited. Interestingly, we did not find that improvement of TCA cycle activity upon adaptive mutation increases the cellular ATP pools. In contrast, we found ATP levels in the adapted strain to be lower than in any other strain, despite the fact that the adapted strain showed a prolonged growth phase and produced more MP1 than the other strains. Accordingly, it is unlikely that increased needs for ATP upon plasmid acquisition were causative for the observed growth deficiency of the original recipient strain RN-T.

It is tempting to speculate that MP1 production entailed excessive channelling of TCA cycle intermediates into the biosynthesis of amino acids, ultimately leading to premature growth arrest of the strain. Along this line, BGC-acquisition had a wide impact on the metabolome of *S. aureus* and adaptive mutation shifted the general pattern a second time to reflect largely that of the plasmid free parental strain. This realization suggested that metabolic fluxes needed to be normalized by improvement of *citZ* expression and the associated increased availability of citrate. Interestingly, deletion of bacteriocin biosynthesis genes in enterococci has been associated with improved growth characteristics and altered expression of ribosomal proteins, vitamins, and glycolysis enzymes (40). Similarly, we found that restoration of the *citZ* allele also entailed a strong increase in expression of the translational machinery, cofactor biosynthesis and of the urea cycle, all suggesting an increase in protein biosynthesis and turnover, supporting the positive effect of the mutation on the cellular metabolism.

The impact of BGC acquisition on TCA cycle activity in *S. aureus* RN4220 is further underlined by the fact that the initial transformant produced high levels of biofilm, which was abrogated upon adaptive mutation in *citZ*. Several studies have associate TCA cycle activity with the staphylococcal capacity to form biofilms. A major factor determining staphylococcal biofilm formation is the production of the PIA (41, 42). We found the responsible *ica* genes to be strongly upregulated upon pD4-19 acquisition. Vuong et al. described that reduced TCA cycle activity increased PIA production in *S. epidermidis* (43), and TCA-deficient mutants were shown to derepress the *ica* genes and to channel carbohydrates into PIA-synthesis (44). We found strongly increased expression of *ica* genes upon pD14-19 acquisition, supporting the general finding that MP1 production induces a TCA deficient phenotype in RN4220. Additionally, it was described that citrate as well as TCA cycle intermediates can stimulate expression of the fibronectin-binding proteins FnbA and FnbB entailing *ica*-independent biofilm formation (45). However, we did not observe differences in *fnbA/fnbB* expression levels in our experiments, suggesting that this mechanism is not relevant for biofilm formation in *S. aureus* RN-T.

Restoration of the *citZ* allele also enhanced MP1 production by the RN4220 lineage. It is well accepted, that expression of antibiotic BGCs in heterologous hosts can be limited due to inappropriate supply of cellular precursors. This is of special relevance for compounds produced by non-ribosomal peptide synthetases or by polyketide synthases which frequently rely on special precursors such as non-proteinogenic amino acids or unusual carbohydrates. For example, overexpression of rhamnose and forosamine biosynthetic pathways improved the biosynthesis of the polyketide antibiotic spinosad 1000-fold (46), and medium optimisation to provide appropriate precursors has proven to be an efficient strategy to enhance compound production (12). Similarly, optimisation of the microbial central metabolic processes including glucose, amino acid, or fatty acid metabolism can boost compound production in heterologous hosts (47). For RiPPs like MP1, the relevance of this concept is less clear, as the compounds are produced using the ribosomal machinery, which relies on the cellular pool of canonical aminoacyl-tRNAs. This might ensure that precursor molecules are generally available, and compound production is possible. However, it seems plausible that RiPP production can drain the pool of aminoacyl-tRNAs. This hypothesis is supported by the finding that addition of the amino acids glutamate, glycine, serine, and threonine as well as the addition of maltose enhanced the production of the RiPP gallidermin (48). However, our metabolomic analysis showed that, in the context of MP1 production, amino acid levels were not depleted in the BGC recipient at the timepoint of growth arrest arguing against amino acid limitation as a sole underlying reason for the decreased compound production.

Finally, it has to be considered that effects that are independent of cellular metabolites might impact the fitness of antibiotic producers. For instance, production of the lantibiotics epidermin and gallidermin imposes a physiological burden on the producing staphylococcal strains because of insufficient immunity of the producer strains leading to increased cell lysis (17). In contrast, we did not observe imperfect resistance of MP1 producing strains, suggesting that insufficient immunity does not substantially contribute to the observed growth defect of RN-T. Most likely the different findings for MP1 and epidermin are due to the different modes of action and the associated resistance mechanisms. Lantibiotics target lipid I, II, III in the bacterial membrane, ultimately damaging the integrity of the cell envelope and inducing lysis (17, 49). Resistance is imperfect and relies on the active expulsion of lantibiotics (50, 51). In contrast, MP1 targets the bacterial ribosome (20) and full resistance is provided by the expression of an alternative L11 ribosomal subunit (52). Interestingly, accumulation of amino acids similar to that observed in the pD4-19 recipient is known to be induced by antibiotic compounds targeting the bacterial ribosome (26). However, expression of the plasmid-associated resistance determinant did not change upon adaptive mutation in *citZ* suggesting that altered levels of autoimmunity are not causative for the observed increase of cellular fitness. However, it seems possible that the malfunctioning *citZ* allele might not be translated in the context of the alternative ribosomal subunit which might explain the TCA-deficient phenotype of the transformant. In general, our observations support a model in which pD4-19 acquisition and associated compound production drain the levels of TCA-cycle intermediates and cause pleiotropic effects including growth deficiency, reduced compound production and increased biofilm formation. These findings enhance our understanding of how antibiotic production is integrated into and optimized by the cellular central metabolism.

Even in environments such as the nasal microbiome bacteriocin producing strains remain rare, although the horizontal transfer of the responsible BGCs is possible and competitive benefits of inhibiting competitors should be immense. Our study can explain why most BGC remain rare among isolates of a given bacterial species. As the consequence of a substantial metabolic burden imposed by bacteriocin production, a BGC-carrying mobile genetic element will most likely be counter selected. Accordingly, The BGCs will only be maintained if adaptive evolution events such as that reported in our study are possible and quickly occurring while simultaneously bacteriocin sensitive competitors are efficiently cleared from the ecological niche. However, further experimental evidence is needed to support this idea.

## Material and Methods

### Strains and growth conditions

The *Staphylococcus* strains used in this study were *S. aureus* D4-19, *S. aureus* RN4220 (RN), and *S. aureus* USA300 LAC. Further strains generated during this study were *S. aureus* RN4220 pD4-19 (RN-T), *S. aureus* RN4220 pD4-19 adapted (RN-A), *S. aureus* D4-19 TN1/2/3, *S. aureus* RN4220 pD4-19 ΔMP1, *S. aureus* D4-19 ΔPP, and *S. aureus* RN4220 pD4-19 ΔPP. Overexpression strains were constructed in *S. aureus* RN4220 background carrying the plasmid pRB473-xylR-6xHis-citZ, with *citZ* deriving from either *S. aureus* RN4220 pD4-19 or *S. aureus* RN4220 pD4-19 adapted. The construction of the plasmids and knockouts is described below. *Escherichia coli* DC10B or *E. coli* Sa08B were used as the cloning host for further transformation in *S. aureus* D4-19, *S. aureus* RN4220 or *S. aureus* RN4220 pD4-19.

Basic medium (BM; 1% soy peptone A3 [Organotechnie SAS, France], 0.5% Ohly Kat yeast extract [Deutsche Hefewerke GmbH, Germany], 0.5% NaCl, 0.1% glucose, and 0.1% K_2_HPO_4_, pH 7.2) was used as the standard growth medium. If necessary, antibiotics were supplemented at a concentration of 10 μg ml^−1^ for chloramphenicol, 2.5 µg ml^-1^ for erythromycin or 0.5 µg ml^-1^ for penicillin G. *E. coli* transformants were grown in lysogeny broth (LB; Lennox) medium (1% tryptone, 0.5% yeast extract, and 0.5% NaCl; Carl Roth GmbH, Germany) supplemented with 10 μg ml^−1^ chloramphenicol or on BM agar with 10 μg ml^−1^ chloramphenicol. BM without glucose (B_0_) was used for expression of *citZ*.

To monitor growth over time, strains were grown overnight in BM under continuous shaking at 37°C. Each strain was adjusted to an optical density at 600 nm (OD_600_) of 1 in BM, and 5 μl of the bacterial stock solutions were pipetted to 1 ml BM into a 24-well microtiter plate. If necessary, 5 mM sodium citrate was added to each well. The plates were incubated for 24 h under continuous shaking in a microplate reader, and OD_600_ was measured every 15 minutes.

### Cloning

DNA manipulation, isolation of plasmid DNA, and transformation of *E. coli* and *S. aureus* were performed by using standard procedures. Enzymes for molecular cloning were obtained from Thermo Fisher Scientific.

### Transposon mutagenesis

To identify the biosynthetic gene cluster responsible for antimicrobial activity of *S. aureus* D4-19, the strain was transformed with the plasmid pBTn and mutants were generated by transposon insertion as described previously (53). To identify the insertion site of the transposon in clones, that had lost antimicrobial activity, genomic DNA was isolated, and an inverse PCR was performed. Therefore, 5 µg gDNA were digested with the restriction enzyme BspHI for 3h at 37°C. After purification of the digest, 2 µg of DNA were religated in 100 µl total volume for 2-3 hours and 2 µl of the ligation mixture were used for standard PCR (in 25 µl volume) with pBTn up and down primers (Table 1). Analyse the PCR products on an analytical gel, isolate strong bands, and sequence with pBTn up/down primers.

**Table 1:**
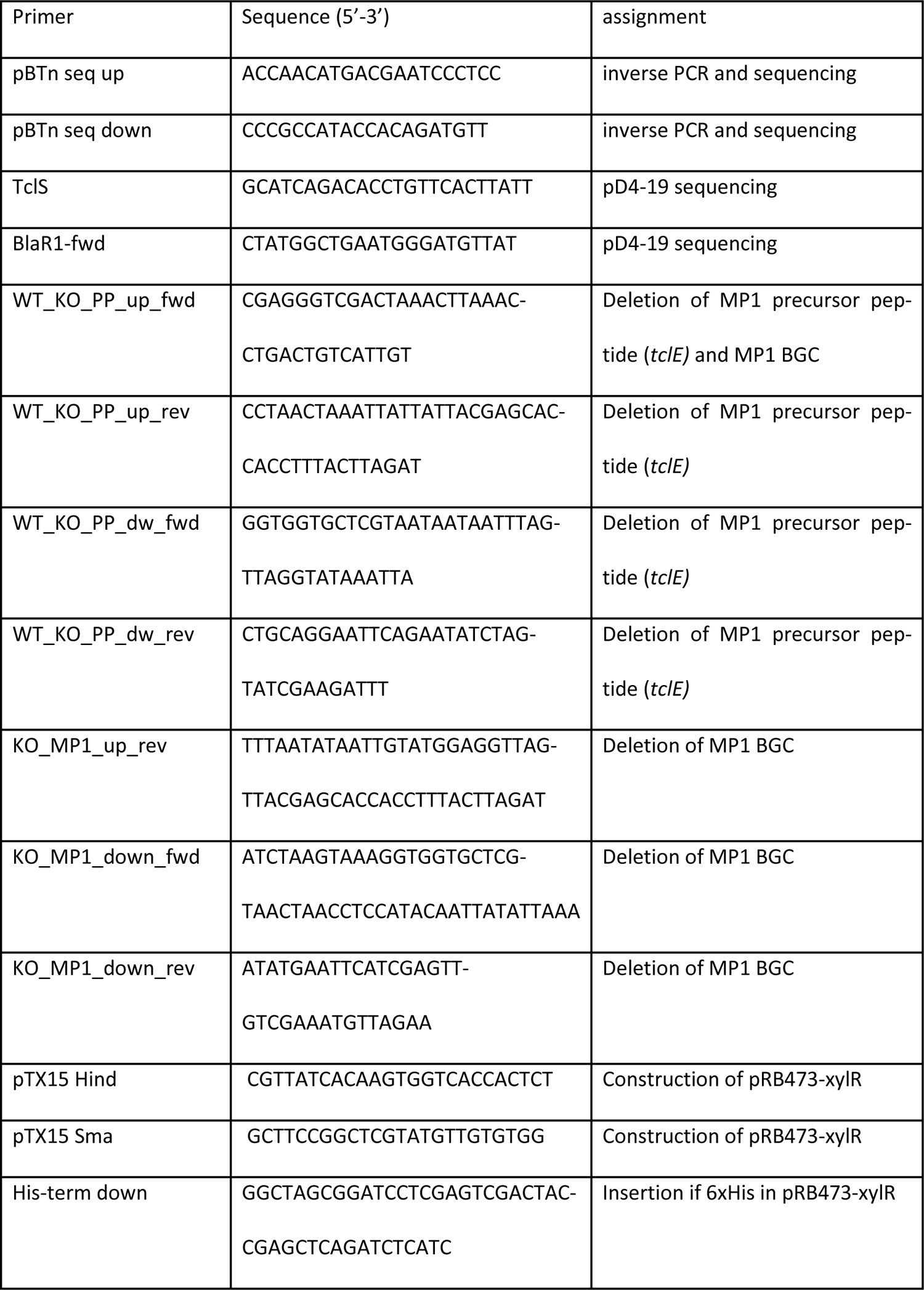
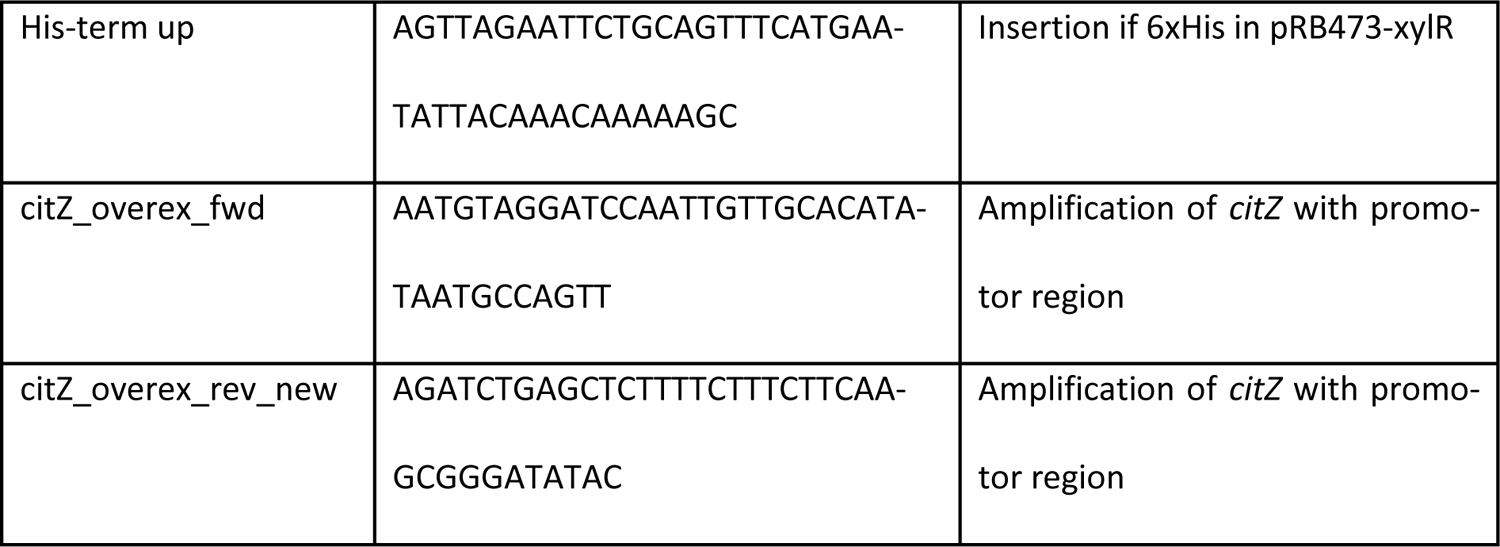
Primers used for transposon mutagenesis and generation of knockout mutants.

### Generation of *S. aureus* RN4220 pD4-19 and adapted mutant

*S. aureus* D4-19 belongs to clonal complex 8, and, to circumvent restriction modification (RM) barriers, we used the laboratory strain *S. aureus* RN4220 as recipient strain. This RM-deficient strain belongs to the same clonal complex, exhibits high transformation rates, and does not show inhibitory effects on other staphylococci. Finally, *S. aureus* RN4220 is highly sensitive to penicillin G (MIC = 0.015 µg/ml) allowing for the selection of pD4-19 which encodes a β-lactamase. Plasmid DNA was isolated from *S. aureus* D4-19 and was used to transform *S. aureus* RN4220 by electroporation. Positive transformants were selected on BM plates containing 0.5 µg ml^-1^ Penicillin G and screened by PCR with primers binding in the BGC and the β-lactamase gene cluster (Table 1). Plasmid isolation and restriction digestion confirmed that the strain *S. aureus* RN4220 had acquired the plasmid pD4-19 and propagated it as an extrachromosomal element.

To generate a mutant that had adapted to MP1 production, *S. aureus* RN4220 pD4-19 was passaged for 28 days by inoculating fresh BM medium every day with 10 µl of overnight culture.

### Generation of knockout mutants

For the generation of knockout mutants, the temperature-sensitive shuttle vector pIMAY was used, and mutants were generated by allelic replacement as described previously (54). Flanking regions of the genes to be deleted were amplified by PCR (Table 1) and ligated into pIMAY after digestion with suitable restriction enzymes. Cloning was performed in *E. coli* DC10B or *E. coli* Sa08B. Sequence-verified plasmids were transferred in the target strains, *S. aureus* D4-19 and *S. aureus* RN4220 pD4-19. Successful knockouts were confirmed by PCR with respective primers and sequencing of the PCR product covering the area of the knocked-out gene.

### Overexpression of *citZ*

To overexpress *citZ*, a novel xylose-inducible plasmid was constructed on the basis of the shuttle vector pRB473. The regulatory unit containing the *xylR* repressor gene and about 200 nucleotides at its 3’-end encompassing the regulated promotor were amplified with primers pTX15 Hind and pTX15 Sma and plasmid pTX15 as template (Table 1). From the resulting PCR product, the required fragment was generated via HindIII and BamHI digestion, which was subsequently ligated to the equally digested pRB473 resulting in pRB473-xylR. Furthermore, a 6xHis tag was inserted into the plasmid to get a C-terminally tagged protein. Therefore, plasmid pPSHG3 was used as template to amplify the 6xHis tag with the primers His-term down and His-term up (Table 1). The insert and pRB473-xylR were digested with EcoRI and BamHI and after ligation, *E. coli* DC10B was transformed. *citZ* was amplified from *S. aureus* RN4220 pD4-19 and *S. aureus* RN4220 pD4-19 adapted using the primers citZ_overex_fwd and citZ_overex_rev_new (Table 1). After restriction digestion with BamHI and SacI, the inserts were ligated into pRB473-xylR-6xHis. Sequence verified plasmids were transferred into *S. aureus* RN4220. To check the expressed protein levels, strains were grown overnight in BM, inoculated to OD_600_=0.1 in B_0_ and grown to OD_600_=0.5. After induction with 0.5% xylose, samples were taken at time points 0 h, 2 h and 4 h. Samples were loaded on an SDS-Page (12% pre-cast gel, BioRad) after incubation with 4x Laemmli Sample Buffer (BioRad, add. 355 mM 2-mercaptoethanol) at 95°C for 5 min and centrifugation for 10 min at 16 000 x*g*. PageRuler Prestained (ThermoScientific) was used, and the gel was run at 120 V for 1 h. For Western Blotting, a nitrocellulose membrane was cut to the size of the gel and the blotting paper, and the membrane were equilibrated in transfer buffer. Blotting paper, gel, membrane, and blotting paper were stacked in a blotting chamber of the BioRad Transblot Turbo. To detect the His-Tag, the membrane was incubated in blocking buffer and then washed with PBS-T (PBS-Tween) and PBS. After incubation with the primary antibody from QIAGEN (Penta-His Antibody, 1:1000 dilution) at RT for 1 h under shaking, the membrane was washed again, 3x with PBS-T and 1 with PBS. The secondary antibody from LICOR (IRDye® 680RD Goat anti-Rabbit IgG, 1:10000 dilution) was added and after incubation for 1h at RT with shaking, washed again as mentioned before. The membrane was analysed using LI-COR Odyssey. Protein amounts were calculated using the LI-COR Odyssey imaging tools.

### MIC assay

Strains used for MIC determinations were grown overnight in BM under continuous shaking at 37°C. Each strain was adjusted to OD_600_=0.00125 in BM and 200 µl were pipetted in the first well of each row of a 96-well microtiter plate. 100 µl were pipetted in the remaining wells, one well per row was used as blank control containing 100 µl BM. The MP1 stock solution was serially diluted in this 96-well microtiter plate and the plates were incubated at 37°C for 21 h under continuous shaking (160 rpm). OD_600_ of cultures in each well was measured with a microplate reader, and the lowest concentration of MP1 leading to no bacterial growth was defined as the MIC value.

### Citrate assay

Citrate levels were measured using the Citrate Assay Kit (MAK057) from Sigma-Aldrich. Strains were grown over night in BM, diluted into fresh BM to OD=0.1 and grown for 5 h at 37°C under continuous shaking. For each strain, 1×10^8^ cells in 100 µl citrate assay buffer were homogenised in a 1.5 ml microcentrifuge tube containing 100 µl glass beads with a Fast prep at 6.5 m/s for 60 s. After centrifugation for 10 min at maximum speed, 30 µl of supernatant were pipetted in a 96 well plate. 20 µl of citrate assay buffer was added to reach the final volume of 50 µl described in the kit manual. Reaction mixes were prepared, and the analysis was carried out as described in the manufacturer’s manual.

### Acetate assay

Acetate levels were measured using the Acetate Colorimetric Assay Kit (MAK086) from Sigma-Aldrich. Strains were grown over night in BM, fresh BM was inoculated to OD=0.1 and grown for 8 h at 37°C under continuous shaking. At time points 0, 2, 4, 6 and 8 hours, 1 ml of culture was centrifuged at 11 000 *xg* for 5 min and the supernatant was transferred into a fresh tube. Samples were 1000-fold diluted, reaction mixes were prepared, and the analysis was carried out as described in the manufacturer’s manual.

### DNA isolation and sequencing

DNA isolation, library preparation and sequencing were performed by the Institute for Medical Microbiology (part of the NGS Competence Center NCCT, Tübingen, Germany). DNA was extracted using the Qiagen Genomic Tip 20/G Kit, following the manufacturer’s instructions. The genomic DNA was quantified with a Qubit dsDNA BR Assay Kit (Thermo Fisher). ONT library preparation was performed following the instructions manual Native barcoding genomic DNA (with EXP-NBD196 and SQK-LSK109, Oxford Nanopore) with an input of 250 ng DNA. 12 µl template DNA was supplemented with the required reagents from the NEBNext Ultra II End Repair/dA Tailing kit (E7546S, NEB) and was first incubated at 20°C for 5 minutes and then at 65°C for 5 minutes. For the barcode ligation 3 µl of nuclease-free water, 0.75 µl End-prepped DNA, 1 µl Native Barcode (Native Barcoding Expansion 96, EXP-NBD196) and 5 µl Blunt/TA Ligase Master Mix (NEB Blunt/TA Ligase Master Mix, M0367) were combined in a new reaction vessel and incubated for 20 minutes at room temperature. 1 μl of 0.5 M EDTA was added and samples were pooled in a new reaction tube. The pool was cleaned-up by using AMPure XP Beads (Agencourt), washed twice with 70% ethanol and resuspended in nuclease-free water. For barcode ligation 5 µl Adapter Mix II, 10 µl NEBNext Quick Ligation Reaction Buffer (5X) and 5 µl Quick T4 DNA Ligase were added to the pool and incubated for 10 minutes at room temperature. The pool was cleaned up using AMPure XP Beads, washed twice with Long Fragment Buffer and eluted in Elution Buffer. The library pool was loaded on a MinION device (Oxford Nanopore Technology, ONT) and stopped at 39 Gb output. Base calling was performed using the ONT’s Guppy basecaller version 4.1.1.

Libraries for Illumina short-read sequencing were prepared using the Illumina Nextera™ DNA Flex Library Preparation Kit with IDT for Illumina DNA/RNA UD indexes, tagmentation according to the manufacturer’s protocol with 500 ng DNA input and 5 cycles indexing PCR. Libraries were checked for correct fragment length on an Agilent 2100 Bioanalyzer and pooled equimolarly and quantified with Qubit DNA HS Assay Kit (ThermoFisher). Equimolarly pooled libraries were sequenced on a MiSeq Reagent Kit v2 (300 cycles) flow cell (Illumina) with 2 x 150 bp read length. For demultiplexing bcl2fastq v2.19.0.316 was used (https://emea.support.illumina.com/downloads/bcl2fastq-conversion-software-v2-20.html).

### DNA data assessment and analysis

Sequencing statistics including the quality per base and adapter content assessment of Illumina reads were conducted with FastQC v0.11.8 (http://www.bioinformatics.babraham.ac.uk/projects/fastqc, accessed June 2022). Unicycler v0.5.0 (55) with default parameters was used for a hybrid assembly of the Oxford Nanopore and Illumina reads of the *S. aureus* RN4220 pD4-19 genome. The resulting genome was annotated using prokka v1.14.6 (56) with the additional parameters to add gene features in the annotation and searching for non-coding RNAs (parameters --addgenes and --rfam). The quality of the assembly was assessed using quast v5.1.0 (57). The genome of the Illumina reads of *S. aureus* RN4220 pD4-19 adapted was assembled using EAGER v1.92.56 (58) and MUSIAL v1.0 (https://github.com/Integrative-Transcriptomics/MUSIAL/tree/v1.0). As a reference the assembly of *S. aureus* RN4220 pD4-19 was used. In EAGER parameters were set to not merge the paired-end reads and to use bwa-mem (59) for the mapping. For SNP calling GATK HaplotypeCaller was chosen (60). Whole-genome sequence of *S. aureus* D4-19 was determined by Illumina short-read sequencing as described above. Illumina reads were *de-novo* assembled with SPAdes (version: 3.9.0) (61) and the plasmid contig was identified with MAUVE (62). The MP1 BGC was analyzed with antiSMASH 5.0 (bacterial settings) (63).

### Spot assay

Antimicrobial activity was assessed, by resuspending the sensitive *S. aureus* USA300 LAC in 200 µl BM and spreading it with a cotton swab on a BM plate. Producer strains were also resuspended in BM and 10 µl of the suspension were spotted on the prepared indicator plate. Once the spots were dry, the plates were incubated at 37°C overnight.

### Purification of MP1

Cell bound MP1 can be isolated as follows. 50 ml overnight culture were centrifuged at 6 000 x*g* for 10 min, the pellet was washed two times with 15 ml PBS and then resuspended in 3 ml methanol. After incubation on a spinning wheel for 1 h and a centrifugation step at 6 000 x*g* for 10 min, the supernatant was transferred in a fresh falcon and used for HPLC or MS/MS analysis.

### LC-MS analysis of MP1

LC-MS analyses were performed with a HPLC (Ultimate 3000, Thermo Fischer) and subsequent HR-ESI(+)-TOF-MS (Maxis 4G, Bruker). For HPLC analysis, LC-MS-grade water (with 0.01 % formic acid) and LC-MS-grade methanol (with 0.06 % formic acid) were used and fractionation was performed with a gradient from 10% to 100% over 20 min and a flow rate of 0.3 ml/min. A nucleoshell RP18 column with a column length of 150 mm, an inner diameter of 2 mm and a particle size of 2.7 µm, prewarmed to 40°C was used. For HR-ESI(+)-TOF-MS sodium formiate was used as calibrant.

### NMR analysis of MP1

NMR analyses were performed on a Bruker AvanceIII-700 instrument. ^1^H NMR spectra were recorded with a frequence of 700 MHz and ^13^C NMR spectra were recorded with a frequence of 176 MHz, both at a temperature of 303 K.

### Sample preparation for metabolome analysis

For the preparation of samples for metabolome analysis, strains were grown over night, inoculated in fresh medium (20 ml) to OD_600_=0.1 and grown for 5 h at 37°C under continuous shaking. OD_600_ was measured and all strains were set to the lowest OD measured. The culture was filtered through a 0.22 µm bottle-top sterile filter (250 ml, Nalgene) via vacuum. The filter was washed with 0.6 % ice-cold NaCl, cut in 4 pieces and incubated at −20°C for 20 min with 5 ml ice-cold 40:40:20 (v/v/v) methanol:acetonitrile:water in a glass bottle. 1 ml of filtrate was transferred into a microcentrifuge tube and bead beaten 2x 30 s at 6.5 m/s with 0.5 ml glass beads. Samples were kept on ice in between. After centrifugation for 5 min at 4°C at maximum speed, 600 µl of supernatant were stored at −80°C.

### Metabolite analysis by flow injection MS (FI-MS)

Metabolites were analyzed by flow injection into a high-resolution quadrupole time-of-flight (QTOF) mass spectrometer (Agilent QTOF 6546) as described previously (64). 3 µL of the sample was injected with an Agilent 1290 Bio Multisampler (G7137A) into the mobile phase that was a 60:40 (v/v-%) mixture of isopropanol (LiChrosolv Supelco hypergrade for LC-MS, 1.02781.2500) and ultrapure water (Omnia Pure, stakpure), buffered with 10 mM ammonium carbonate ((NH_4_)_2_CO_3_, Sigma-Aldrich, 3799-10 G) and 0.04% ammonium hydroxide (NH_4_OH, Honeywell/Fluka TraceSELECT Ultra, 16748-250 ML). The flow rate of the mobile phase was 0.15 ml*min^-1^. Mass spectra were separately recorded in positive- and negative ionization profile mode from *m/z* 50 to *m/z* 1700 with an acquisition rate of 1.4 ms/spectrum using the highest resolving power (10 GHz-High Sensitivity). Online mass axis correction was performed with purine and hexakis(1H,1H,3H-tetrafluoropropoxy)phosphazine (HP-0921, Agilent Technologies). The source gas temperature of the ESI ion source was 225 °C, with 11 l*min^−1^ drying gas and a nebulizer pressure of 20 psi. The sheath gas temperature was 350 °C, and the flow rate was 10 l*min^−1^. Electrospray nozzle and capillary voltages were 2,000 and 3,500 V, respectively. Fragmenter and skimmer voltages were 120 and 65 V, respectively. Ion peaks were annotated by matching the mass-to-charge ratios to calculated, single (de-) protonated masses of metabolites listed in a genome scale model of *Escherichia coli* K12 (iML1515) (65). Hierarchical cluster plot was generated using MATLAB. Metabolites obtained via FI-MS, which were used for further analysis, are listed in table 2.

**Table 2:** Metabolites and their isobars measured via FI-MS.

| metabolite(s) | m/z |
| --- | --- |
| citrate/isocitrate/5-dehydro-4-deoxy-D-glucarate [M-H]<sup>-</sup> | 191.020 |
| α-ketoglutarate [M-H]<sup>-</sup> | 145.014 |
| malate [M-H]<sup>-</sup> | 133.014 |
| fumarate [M-H]<sup>-</sup> | 115.004 |
| pyruvate [M-H]<sup>-</sup> | 87.009 |
| lactate/glyceraldehyde/dihydroxyacetone [M-H]<sup>-</sup> | 89.024 |
| acetate [M-H]<sup>-</sup> | 59.014 |
| glucose, galactose, allose, fructose, mannose and myo-inositol [M-H]<sup>-</sup> | 179.056 |
| citrulline [M+H]<sup>+</sup> | 176.103 |
| butyrate [M+H]<sup>+</sup> | 89.060 |
| carnitine [M+H]<sup>+</sup> | 162.112 |
| choline [M+H]<sup>+</sup> | 104.107 |

### Metabolite analysis by LC-MS/MS

LC-MS/MS was performed with an Agilent 6495 triple quadrupole mass spectrometer (Agilent Technologies) as described previously (66). An Agilent 1290 Infinity II UHPLC system (Agilent Technologies) was used for liquid chromatography using two columns: i) an Acquity UPLC BEH Amide (Waters) for acidic conditions and ii) an iHILIC-Fusion(P) (HILICON AB) for basic conditions. The column oven was at 30°C. LC solvents were: solvent A: water with ammonium formate (10 mM) and formic acid (0.1 % v/v) for acidic conditions, and water with ammonium carbonate (10 mM) and ammonium hydroxide (0.2 %) for basic conditions; solvents B: acetonitrile with formic acid (0.1% v/v) for acidic, and acetonitrile for basic conditions. The LC gradient was: 0 min 90% B, 1.3 min 40% B, 1.5 min 40% B, 1.7 min 90% B, 2 min 90 % B. The flow rate was 0.4 ml/min. The injection volume was 3 µL. Settings of the ESI source were: 200 °C source gas, 14 L/min drying gas and 24 psi nebulizer pressure. The sheath gas temperature was at 300 °C and flow at 11 L/min. The electrospray nozzle was set to 500 V and capillary voltage to 2500 V. All samples were mixed with a 13-C labelled internal standard and the ratio of 12-C and 13-C peak heights was used to quantify metabolites. 12C/13C ratios were normalized to the OD-specific cell volume at the time point of sampling. Alanine, arginine, asparagine, aspartate, glutamate, glutamine, glycine, histidine, leucine/isoleucine, lysine, methionine, phenylalanine, proline, serine, threonine, tyrosine, tryptophan, valine, acetyl-CoA, succinate, 3-phosphoglycerate, phosphoenolpyruvate, urea, ornithine, argininosuccinate and ATP were measured via LC-MS/MS.

### Sample preparation for proteome analysis

Strains were grown over night, inoculated in fresh medium to OD_600_=0.1 and grown for 5 h at 37°C under continuous shaking. 1.5 ml were centrifuged, the pellet was resuspended in 1 ml SDS buffer (4% w/v sodium dodecyl sulfate (SDS), in 100 mM tris(hydroxymethyl)aminomethane (Tris)/HCl; pH 8). Cells were homogenised at 6.5 m/s for 2×40s by using a Fast Prep, and incubated on ice for 2 min in between. Samples were centrifuged at maximum speed for 1 min and the supernatant was transferred into a fresh Eppendorf tube. To reduce cysteine disulfide bonds, 10 mM Dithiothreitol (DTT) was added to the samples and incubated for 45 min shaking at 650 rpm at room temperature (RT). 5.5 mM Iodoacetamide (IAA) were added to alkylate reduced cysteine thiol groups. Incubation for 45 min at RT with shaking at 650 rpm in the dark were followed by centrifugation of the samples at 12 000 *xg* for 15 min. The supernatant was transferred in a new tube and one volume of supernatant was mixed with 7 volumes of ice cold 8:1 acetone:methanol, vortexed and incubated over night at −20°C. Centrifugation of the precipitated proteins for 5 min at 1 000 x*g* was followed by two washing steps with 80% acetone at RT. The protein pellet was air dried for 10-15 min and rehydrated in denaturation buffer (6 M urea, 2 M thiourea in 10 mM Tris/HCl; pH 7.5).

### LC-MS/MS analysis of proteome samples

Ten microgram of proteins per sample were digested in solution with trypsin as described previously (67). Desalted peptides (68) were separated on an Easy-nLC 1200 system coupled to a quadrupole Orbitrap Exploris 480 mass spectrometer (all Thermo Fisher Scientific) as described previously (69) with slight modifications: peptides were separated using an 87-minute segmented gradient from 10-33-50-90% of HPLC solvent B (80% acetonitrile in 0.1% formic acid) in HPLC solvent A (0.1% formic acid) at a flow rate of 200 nl/min. The mass spectrometer was operated in data-dependent mode, collecting MS spectra in the Orbitrap mass analyser (60,000 resolution, 300-1750 *m/z* range) with an automatic gain control (AGC) set to standard and a maximum ion injection time set to automatic. The 20 most intense precursor ions were sequentially fragmented with a normalized collision energy of 28 in each scan cycle using higher energy collisional dissociation (HCD) fragmentation. In all measurements, sequenced precursor masses were excluded from further selection for 30 s. MS/MS spectra were recorded with a resolution of 15,000, whereby fill time was set to automatic. Acquired MS spectra were processed with MaxQuant software package version 1.6.14.0 (70) with integrated Andromeda search engine (71). Database search was performed against a *Staphylococcus aureus* (allStrains) protein database (downloaded on 7th of October 2020, 216,059 entries), and 286 commonly observed contaminants. Endoprotease trypsin was defined as protease with a maximum of two missed cleavages. Oxidation of methionine, and protein N-terminal acetylation were specified as variable modifications. Carbamidomethylation on cysteine was set as fixed modification. Initial maximum allowed mass tolerance was set to 4.5 parts per million (ppm) for precursor ions and 20 ppm for fragment ions. Peptide, protein and modification site identifications were reported at a false discovery rate (FDR) of 0.01, estimated by the target-decoy approach (72). The iBAQ (Intensity Based Absolute Quantification) and LFQ (Label-Free Quantification) algorithms were enabled, as was the “match between runs” option (73, 74).

### RNA isolation for transcriptome analysis

Strains were grown over night, inoculated in fresh medium to OD_600_=0.1 and grown for 5 h at 37°C under continuous shaking. 1/10 volume EtOH/Phenol was added to 500 µl of sample and mixed for 1 min. After incubation on ice for 5 min, the samples were centrifuged for 1 min at 20 000 x*g* at 4°C. The supernatant was discarded, and the pellet was resuspended in 1 ml TRIzol. Each sample was transferred to one screw-cap tube with glass beads and cells were lysed via bead-beating for 2x 6.5 ms/s for 30 s. In between the two runs the cells were kept on ice for 2 min. 200 µl chloroform were added to the samples, mixed and incubated 2-3 min before centrifugation of the samples for 15 min, 12 000 x*g*, 4°C. The aqueous supernatant was taken, mixed with 500 µl isopropanol and samples were centrifuged for 10 min, 21 000 x*g*, 4°C. The supernatant was discarded, and the pellet was resuspended in 500 µl 75% EtOH. After centrifugation for 5 min, 20 000 x*g*, 4°C, the supernatant was discarded, and the pellet was dried at RT. The pellet was resuspended in 100 µl RNA-grade water and RNA was concentrated via the MN RNA clean up kit (Machery-Nagel) and RNA was eluted in 60 µl RNA-grade water. After quantification with a Nanodrop, RNA was stored at −80°C. Library Prep and Sequencing was performed by the Institute for Medical Microbiology (part of the NGS Competence Center NCCT (Tübingen, Germany)). RNA samples were DNase I digested (DNase I recombinant, Rnase-free, Millipore Sigma), cleaned up (RNA Clean & Concentrator-5, Zymo Research), quantified (Qubit RNA BR Assay Kit, ThermoFisher) and normalized to 100 ng in 11 µl nuclease-free water. Library preparation was performed according to the Illumina Stranded Total RNA Prep, Ligation with Ribo-Zero Plus Reference Guide. Library concentration was measured with Qubit DNA HS Assay Kit, (ThermoFisher) on a Qubit Fluorometer (invitrogen) and fragment length was assessed with an Agilent 2100 Bioanalyzer (Agilent High sensitivity DNA Kit, Agilent). Samples were equimolarly pooled and sequenced on NextSeq™ 500 High Output Kit v2.5 (75 cycles) flow cell (Illumina) with 1 x 75 bp read length.

### RNA-Seq data assessment and analysis

Sequencing statistics including the quality per base and adapter content assessment of resulting transcriptome sequencing data were conducted with FastQC v0.11.8 (http://www.bioinformatics.babraham.ac.uk/projects/fastqc, accessed June 2022). All reads mappings were performed against the previously assembled reference strain *S. aureus* RN4220 pD4-19 (SRA Bioproject ID PRJNA855446). The mappings of all samples were conducted with HISAT2 v2.1.0 (75). As parameters spliced alignment of reads was disabled and strand-specific information was set to reverse complemented (HISAT2 parameter --no-spliced-alignment and --rna-strandness “R”). The resulting mapping files in SAM format were converted to BAM format using SAMtools v1.9 (76). Mapping statistics, including strand specificity estimation and percentage of mapped reads, were conducted with the RNA-Seq module of QualiMap2 v2.2.2-a (77). Gene counts for all samples were computed with featureCounts v1.6.4 (78) based on the prokka annotation of the assembled *S. aureus* RN4220 pD4-19 genome, where the selected feature type was set to transcript records (featureCounts parameter -t transcript). A quality check for ribosomal rRNA was performed with a self-written script based on the absolute counts of annotated rRNAs. To assess variability of the replicates of each time series, a principal component analysis (PCA) was conducted with the DESeq2 package v1.28.1 (79).

### Normalization and differential gene expression

For the computation of genes differentially expressed between the two different strains (*S. aureus* RN4220 pD4-19 and pD4-19 adapted) and the wild type strain *S. aureus* RN4220, DESeq2 v1.20.0 (79) was applied to the absolute gene counts as computed with featureCounts. For differences between the two strains and the wildtype strain, genes with an adjusted p-value (FDR) < 0.05 and absolute log2 fold change (FC) > 1 were reported as differentially expressed.

### Gene set enrichment analysis

The assembled *S. aureus* RN4220 pD4-19 genome was functionally annotated using FACoP (http://facop.molgenrug.nl/). Gene set enrichment analysis was performed on differentially expressed genes using FUNAGE-Pro (http://gseapro.molgenrug.nl/).

### Biofilm assay

Strains were grown over night in tryptic soy broth and adjusted to the OD_600_ of the strain with the lowest OD_600_. 5 μl were added to 995 μl of BM containing 1% (v/v) glucose. 200 μl were transferred into wells of a fibrinogen coated Nunclon Delta surface microtiter plate. The plate was incubated at 37°C for 24 h, in non-shaking conditions. Control wells with broth and no bacteria were included. The supernatant was discarded, and wells were washed three times with 200 μl PBS. The plate was inverted for 30 min to dry. 100 μl crystal violet was added to each well and washed off after 1 min (3-5 times with PBS). 100 μl of 5% acetic acid was added to the wells and the plate was placed on a shaker for 5 min to dissolve the cells. Subsequently the absorbance at 570 nm was measured using the FLUOstar Optima.

### Statistical analysis

Statistical analyses were performed using GraphPad Prism 9.02.

### Data availability

ONT and Illumina reads for *S. aureus* RN4220 pD4-19 and *S. aureus* RN4220 pD4-19 adapted can be found at SRA Bioproject ID PRJNA855446 and the plasmid sequence of pD4-19 (from *S. aureus* D4-19) has the GenBank accession number ON936820. All RNA-Seq Illumina read files as well as the raw counts have been deposited in NCBI’s Gene Expression Omnibus and are accessible under accession number GSE208001. Metabolome data can be accessed at Metabolights via the identifier MTBLS5196. Proteome data can be accessed at PRIDE via the accession number PXD035193.

## Acknowledgements

The authors thank Libera Lo Presti for critically reading and editing this manuscript and the NCCT (NGS Competence Center Tübingen, Germany) for whole genome and transcriptome sequencing of our strains. Furthermore, we authors thank the Quantitative Biology Center of the University of Tübingen for project and data management.

Our work was funded by the Deutsche Forschungsgemeinschaft (DFG, German Research Foundation) in the frame of the TRR261 project ID 398967434 (S.H., A.P., S.G., K.N., H.L., B.M.), Germanýs Excellence Strategy – EXC 2124 – 390838134 (S.H., A.P., B.K., H.L.), and the TRR156 (A.P.). Further we acknowledge support by the High Performance and Cloud Computing Group at the Zentrum für Datenverarbeitung of the University of Tübingen, the state of Baden-Württemberg through bwHPC and DFG grant no INST 37/935-1 FUGG. Finally, we acknowledge funding by the German Center of Infection Research (DZIF) to S.H., A.P.

